# First endangered black-footed ferrets, *Mustela nigripes*, cloned for genetic rescue

**DOI:** 10.1101/2024.04.17.589896

**Authors:** Ben J. Novak, Pete Gober, Robyn Bortner, Della Garelle, Mary Wright, Justin Chuven, Marlys L. Houck, Oliver A. Ryder, Dennis Milutinovich, Jill Benavidez, Kerry Ryan, Shawn Walker, Sanaz Sadeghieh Arenivas, Lauren Aston, Blake Russell, Paul Marinari, Adrienne Crosier, Kelly Helmick, Mary R. Gibson, Daniel P. Carlsen, Bradley J. Swanson, Samantha M. Wisely, Zoe S. White, Colleen Lynch, Ryan Phelan

**Affiliations:** Revive & Restore, 1505 Bridgeway #203, Sausalito, CA 94965, USA; US Fish & Wildlife Service National Black-footed Ferret Conservation Center, Carr, CO 80612; Beckman Center for Conservation Research, San Diego Zoo Wildlife Alliance, Escondido, CA, 92027, USA; ViaGen Pets & Equine, Cedar Park, Texas, 78613, USA; Smithsonian Conservation Biology Institute, Front Royal, Virginia, USA; Department of Biology, Central Michigan University, Mount Pleasant, MI, USA; Department of Wildlife Ecology and Conservation, University of Florida, 110 Newins-Ziegler Hall, Gainesville, Florida 32611-0430, USA; Population Management Center, Lincoln Park Zoo, 2001 North Clark St. Chicago, 60614

## Abstract

An endangered black-footed ferret female that died in 1988 with no living descendants in the current population was successfully cloned from cryopreserved cells using cross-species somatic cell nuclear transfer, producing three healthy kits. Incorporating progeny from these clones would provide an 8th founder to the breeding program and increase genetic variation to the species’ limited gene-pool. This marks the first time a native U.S. endangered species has been cloned.

## Main Text

The black-footed ferret, *Mustela nigripes*, is one of the most endangered North American mammals. Twice presumed extinct, the species was saved by an intensive captive breeding and reintroduction program^1^. The last few surviving wild individuals were captured between 1985 and 1987 to establish the founding conservation breeding population from which the living descendant population is entirely derived^2^ (Fig. 1). Over 11,000 ferrets have been born since, of which ∼4,600 have been reintroduced to establish 34 wild populations^3^, all of which are descended from just seven individual founders^2^. Careful management has worked to conserve as much founding genetic diversity as possible, however differential reproductive output of the founders compounded by subsequent genetic drift has resulted in a significant loss of variation. As of 2019, studbook pedigree analysis estimates that black-footed ferrets’ mean kinships, ranging 0.1293-0.1496^2^, are slightly higher than that of first cousins (0.125).

**Figure 1.**
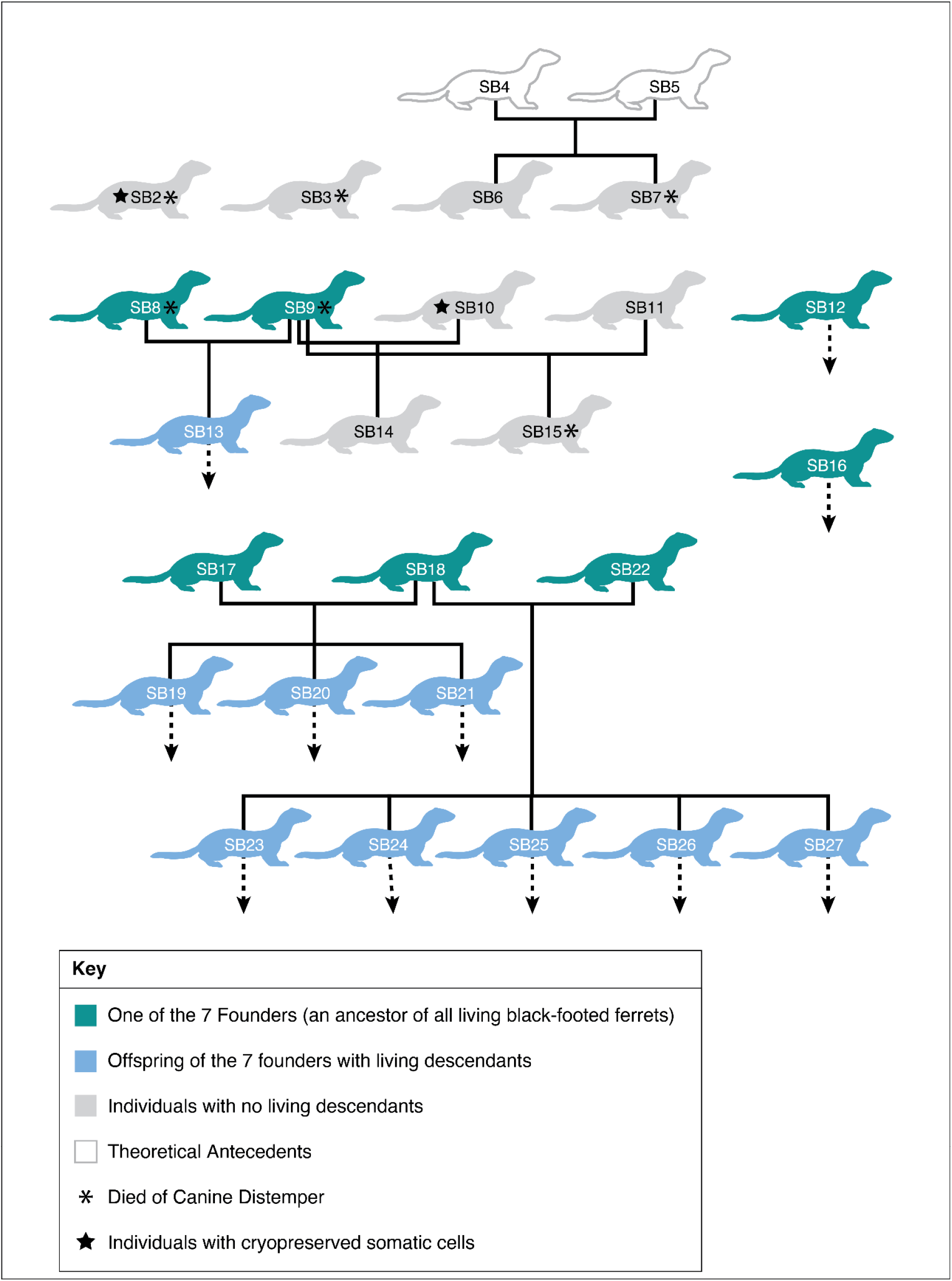
Pedigree of wild-caught black-footed ferrets captured from 1985-1987 to establish the captive breeding program from which all living individuals are descended. Dotted line arrows indicate individuals with living descendants.

While the species is not yet significantly compromised from inbreeding, declines in reproductive fitness correlating with increased inbreeding coefficients have been observed^4^; therefore, long-term viability and recovery can likely be enhanced by genetic rescue^1^. Genetic rescue in practice is the introduction of new, genetically different individuals to a population with low genetic diversity, which when interbred initially reduces the genetic load of the subsequent generations thereby improving the overall fitness of the target population^5^. Cryopreservation of gametes and cells for use with advanced reproductive technologies expands the scope of genetic rescue to include historic individuals as sources of new genetic variation (variation lost during population bottlenecks or to subsequent genetic drift and then restored is “new” variation to present-day populations). In previous genetic rescue attempted in this species, the “new” source of the variation was from cryopreserved sperm collected from founder and early pedigree males who were born 20 generations prior to the females that were impregnated via artificial insemination^6,7^. Although genetic diversity did increase in the F1 generation, an increase in individual fitness was not observed. Because of the genetic similarity and high relatedness of all extant black-footed ferrets, it was recognized that a source of unique individuals outside the current population was needed.

Fortuitously, forward-thinking conservationists cryopreserved somatic cell lines of two historic black-footed ferrets that were captured for *ex situ* breeding, but today have no living descendants (Figure 1), creating the opportunity to introduce new founders to the breeding program via somatic cell nuclear transfer (SCNT)^1^. Here we report the generation of clones from one of those cell lines, Studbook Number 10 (SB10), a female named Willa, using cross-species SCNT. Comparative genomics of the two historic ferret cell lines and more than 50 individuals representing the history of the captive breeding program as well as two reintroduced populations have shown that the historic cell lines possess many times more private alleles than the average living black-footed ferret (unpublished results). Through cloning, it is hoped this historic genetic variation can be reintroduced to benefit the species.

In 2020, a total of 41 cloned embryos derived from adult SB10 donor cells were transferred to two surrogate domestic ferret females, resulting in one successful pregnancy with one live birth and one stillbirth (Table 1). The live clone born December 10, 2020, at the National Black-footed Ferret Conservation Center (NBFFCC) was designated Studbook Number 9819 and named Elizabeth Ann. In 2023, a total of 122 cloned embryos generated from rejuvenated fetal fibroblast donor cells derived from a terminated SB10 cloned fetus were transferred to four domestic ferret surrogate females. Two successful pregnancies resulted, producing two live births and one stillborn kit (Table 1). These live clones, born May 1 at the NBFFCC and May 23, 2023 at the Smithsonian Conservation Biology Institute (SCBI) are designated as Studbook Numbers SB10498 and SB10602, named Noreen and Antonia, respectively. Clones were compared to donor genotype profiles via microsatellite sequencing; all clones are an exact nuclear microsatellite match to the donor (Table 4). PCR, TA TOPO cloning, and Sanger sequencing of partial nuclear and mitochondrial gene sequences confirmed the kits as the result of cross-species SCNT; the kits possess black-footed ferret nuclear DNA and domestic ferret mtDNA from the oocyte donor. Noreen and Antonia are homoplasmic for domestic ferret mtDNA, while Elizabeth Ann exhibits heteroplasmy (20% black-footed ferret mtDNA and 80% domestic ferret mtDNA, Table 5).

**Table 1.** Results of embryo transfers derived from natural fertilization, same-species SCNT, and cross-species SCNT.

| Somatic Cell Donor Species & Cell Line | Donor Cell Type | Embryos Transferred | Recipients | Pregnancies | Number of Fetuses at 28 Days Gestation (% Embryos Transferred) | Live Births (% Embryos Transferred) |
| --- | --- | --- | --- | --- | --- | --- |
| *Naturally conceived M. putorius furo* embryos | NA | 11 | 1 | 1 (100%) | 6 (54.5%) | NA |
| *M. putorius furo* | Fetal fibroblasts | 26 | 1 | 1 (100%) | 1 (3.8%) | NA |
| *Mustela putorius furo* | Adult fibroblasts | 42 | 1 | 1 (100%) | 1 (2.4%) | NA |
| *M. nigripes* SB8351 | Adult fibroblasts | 86 | 3 | 2 (66%) | 5 (5.8%) | NA |
| *M. nigripes* SB10 | Adult fibroblasts | 41 | 2 | 1 (50%) | 2 (4.9%) | 1 (2.4%) |
| *M. nigripes* SB10 | Rejuvenated fetal fibroblasts | 122 | 4 | 2 (50%) | 3 (2.4%) | 2 (1.6%) |

All living clones grew normally (Figure 2), reaching developmental and behavioral milestones at the times expected for naturally conceived black-footed ferrets^8^ (Table 2), and continue to be in good health at the time of writing this report. The clones born in 2023 will not reach sexual maturity until March-July 2024. Elizabeth Ann came into estrus during the 2022 breeding season at 15 months of age (female black-footed ferrets usually reach sexual maturity at ∼10-12 months of age, but Elizabeth Ann was born outside the natural breeding season, hence her older age of first estrus). She rejected the males presented for natural mating, a not uncommon occurrence in *ex situ* breeding programs. Elizabeth Ann was prepped for artificial insemination, at which time she was found to have a pathologic uterine condition, hydrometra and non-infectious metritis, and therefore she was not inseminated. Her uterine swelling progressed in severity necessitating an ovariohysterectomy to prevent the condition from becoming fatal, thus precluding future breeding. Elizabeth Ann recovered well and was healthy at the time of writing this report at 37 months of age.

**Table 2.** Comparison of cloned black-footed ferret developmental milestone timelines to the average age of developmental milestones in naturally conceived black-footed ferrets, as previously compiled by^8^.

| Trait | Average Day of First appearance | Observed for Elizabeth Ann | Observed for Noreen | Observed for Antonia |
| --- | --- | --- | --- | --- |
| Face mask pigmentation | 16 | 16 | 15 | 13 |
| Begin eating solid food | 32 | 25 | 35 | 34 |
| Ears unfold | 32 | 32 | 31 |  |
| Eyes open | 37 | 36 | 36 | 38 |
| Vocal repertoire expands | 37 | 38 | 45* | 44* |
| Wandering on cage surface | 49 | 43 | 50 | 46 |
| Permanent canines and molars erupting | 63 | 60 | 60 |  |
| Observed killing live prey | 70 | 84** | 85** | 95** |
| Complete adult dentition | 84 | 84 | 84 |  |

\*This was the day the vocal repertoire expansion was first noted, but this may have started earlier.
\*Prey was not offered earlier to the clones. The clones killed live prey when it was first offered to them.

**Figure 2.**
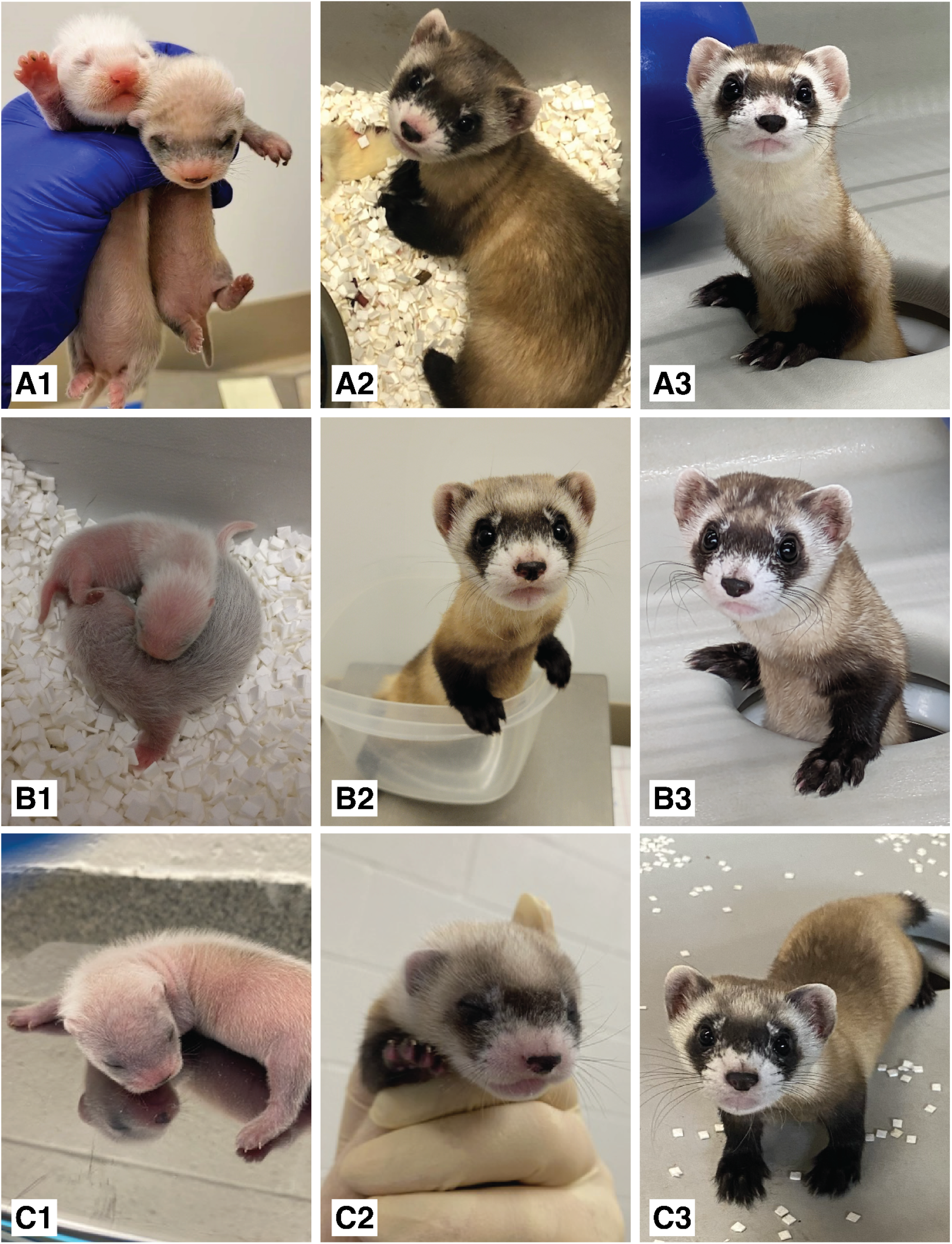
Images of all three cloned black-footed ferrets at intermittent life stages. A1-A3: Elizabeth Ann at 16 days (pictured with domestic ferret foster litter mate), 48 days, and one year of age. B1-B3: Noreen at 10 days (pictured with domestic foster litter mate), 52 days, and 136 days of age. C1-C3: Antonia at 13 days, 29 days, and 70 days of age.

Elizabeth Ann had just one fully developed uterine horn and only one kidney (both on the right-hand side). A unilateral renal/reproductive system phenotype has been observed in several naturally conceived female black-footed ferrets, and a single kidney has also been observed in naturally conceived male black-footed ferrets. At least one naturally conceived black-footed ferret female with a single uterine horn has produced healthy offspring previously, thus the phenotype does not appear to prevent reproduction. There is no evidence that this phenotype is correlated to inbreeding, though a thorough analysis has not been performed. No records of SB10’s internal anatomy could be found for comparison, but the stillborn clone born in 2023, generated by rejuvenated fetal fibroblasts, was confirmed during necropsy to have two kidneys and two uterine horns. The two live clones born in 2023 from rejuvenated fetal fibroblasts, Noreen and Antonia, also have two kidneys each as found by physical palpation during their 60-day medical exams, but the existence of two uterine horns cannot be ascertained by a physical examination and at this time is still undetermined; due to the close developmental relationship between the reproductive and renal systems we expect that Noreen and Antonia possess two uterine horns. We infer that the unilateral renal/reproductive system of Elizabeth Ann is not a genetic trait, but the result of some epigenetic or environmental factor during development. Noreen and Antonia, offer another chance to breed this genetic line and become new founders for the population when they reach sexual maturity in spring 2024.

By the International Union for the Conservation of Nature (IUCN) assessment and classifications of endangerment (Near Threatened, Vulnerable, Endangered, Critically Endangered, Extinct in the Wild), the black-footed ferret is now the sixth endangered species to be successfully cloned^9^ and the first U.S. native endangered species to be cloned. More significantly, this is the first time that cloning has been used to produce new, genetically unrelated individuals for a population that has experienced a genetic bottleneck. This work showcases the value of biobanking and advanced reproductive technologies for other declining and recently bottlenecked species for which survival depends on careful genetic management^10-12^. For black-footed ferrets, this breakthrough establishes the opportunity to use additional cryopreserved materials and cloning as a means to combat genetic drift and inbreeding depression and ensure genetic viability within the conservation breeding program for as long as the program is needed to achieve, secure, and maintain the recovery of the species. Without the genetic rescue afforded by biotechnology, the genetic diversity of this closed population would otherwise steadily decline^1^, continuing to compromise fitness and reproductive success, potentially rendering the species conservation reliant even with successful habitat restoration and management. These clones and future genetic rescue work built upon the foundations of cloning^1,13^ offer a future in which independent recovery for the species is attainable.

## Methods

All work performed was carried out under the conditions established by the Black-footed Ferret Managed Care Operations Manual^14^.

### The Donor Individual: SB10, Willa

A post-mortem skin biopsy from a female black-footed ferret (IDW391) was received at the San Diego Zoo Center for Reproduction of Endangered Species from the Wyoming Department of Fish & Game in January 1988. This female was captured from the wild on November 2, 1985 and assigned Studbook number 10 (SB10) in the breeding pedigree. Cell culturing, harvesting, and chromosome banding followed the techniques described by Kumamoto et al^15^. Briefly, the skin biopsy was processed for fibroblast cell culture by explanting tissue fragments onto the base of a T25 flask. Cells were grown and expanded at 37°C in media enriched with 10% fetal bovine serum, 1% glutamine and 1% penicillin/streptomycin and frozen in liquid nitrogen with 10% DMSO at low passage and stored long-term at the San Diego Zoo Wildlife Alliance Frozen Zoo® with a lab identifier of KB6253. SB10 was bred but did not produce offspring. A wild-caught male presumed to be her son, SB14, also did not sire offspring, terminating the lineage and resulting in SB10 having no living descendants.

### Cloning Process

To clone black-footed ferrets, cross-species SCNT was conducted by ViaGen Pets & Equine, following previously established methods for same-species SCNT of domestic ferrets, *Mustela putorius furo* ^16,17^. Cross-species SCNT was implemented to avoid the use of rare black-footed ferret females as oocyte donors and recipient surrogate mothers, allowing female black-footed ferrets to maximize their reproductive output through natural breeding to maintain conservation recovery efforts. Oocytes for embryo reconstruction were obtained from domestic ferrets post-mortem (these ferrets were sacrificed for other research purposes, not for the purpose of cloning black-footed ferrets). Cloned embryos were reconstructed via electrofusion, introducing the entire cytoplasmic contents of the black-footed ferret donor cells into the domestic ferret oocytes.

A freshly established, non-cryopreserved fibroblast cell line from a living male, SB8351, was used for preliminary cloning trials to evaluate cross-species nuclear-cytoplasmic and gestational compatibility with domestic ferret oocytes and surrogate mothers. A total of 86 cloned embryos of SB8351 were transferred to three surrogates, resulting in two pregnancies that produced one and four fetuses respectively (Table 1). Cloned fetuses were collected at day 28 of the 42-day gestation and compared to domestic ferret fetuses of the same developmental stage that were produced via natural fertilization followed by embryo transfer and same-species SCNT using adult and fetal fibroblast donor cells (Table 1). Microsatellite genotyping (Table 4) and Sanger sequencing of diagnostic partial gene sequences of the black-footed ferret fetuses confirmed each clone as a genetic match to the original somatic donor with a mixture of both black-footed ferret and domestic ferret mitochondrial genomes. Black-footed ferret cloned fetuses were developmentally normal, indicating cross-species compatibility and prompting full-term cloning trials using the SB10 cell lines. After the birth of the first clone from adult donor cells, further trials used rejuvenated fetal fibroblasts to produce additional clones. In both 2020 and 2023 the cloned embryos were generated and transferred to recipient surrogates by ViaGen Pets & Equine. Pregnancies verified by ultrasound between days 15-25 of gestation were transported to the NBFFCC and SCBI at day 21-28 gestation to complete gestation to term (42 days).

### Morphological and Behavioral Development

Elizabeth Ann, Noreen, and Antonia, although generated using domestic ferret oocytes, gestated in domestic ferret uterine environments, and raised by a domestic ferret foster families including domestic ferret litter mates, exhibited normal morphological and behavioral development for black-footed ferrets^8^. The standard milestones were observed and noted throughout the development of the clones, as well as observations of other noteworthy behavioral traits.

While Elizabeth Ann was submissive and docile for the first few weeks of life, caretakers noted more aggressive and playful behavior upon opening her eyes for the first time (38 days old), similar to her domestic litter mates and black-footed ferret kits of the same age. After weaning and into adulthood, Elizabeth Ann has exhibited a “middle ground” behavior for black-footed ferrets between that of an older docile individual and the most aggressive or timid individuals. During the observation period, she exhibited the typical elusiveness of black-footed ferrets, spending the majority of her time in the burrow, or “underground”, portion of her housing (the housing is comprised of tubing that leads to a sublevel darker den underneath the “surface” level open-light housing unit). She came to the “surface” primarily to feed. She played with and attacked objects when presented with them and successfully killed live prey (Table 2). When caretakers have opened the housing unit and attempted to shift her to a handling cage for routine health care procedures she reacted defensively and aggressively with posture and vocalization like most other black-footed ferrets. Although currently only an *n* of 3, these observations do support a strong genetic component to black-footed ferret behavior, which bodes well for the continued use of cloning for conservation management in this species.

### Nuclear DNA genotyping of cloned black-footed ferrets by microsatellite sequencing

To validate that cloned fetuses, stillborns, and live clones produced in 2020 were the result of cross-species SCNT, PCR DNA sequences were compared from two nuclear genes and one mtDNA gene that are diagnostic between species. Cross-species clones were expected to possess nuclear DNA from the donor species and mtDNA from the oocyte donor species.

The nuclear DNA sequences amplified were Exon 9 of the Breast and Ovarian Cancer Susceptibility gene (BRCA-1)^18^, and an intron-exon region of the Growth Hormone Receptor gene (GHR)^19^. These sequences are diagnostic between species. The mtDNA gene amplified was Cytochrome b (Cytb)^20^.

Blood samples for genotyping were obtained from live clones during their 60 days-of-age medical exam. A leg was removed from each stillborn clone for DNA extractions before necropsy. Fetal fibroblast lines were derived from terminated clones for genotyping.

DNA was extracted from cell and fetal tissue samples using the Macherey-Nagel Nucleospin Tissue Kit, following published protocols (Machery-Nagel, Düren, Germany). Samples were extracted singly to reduce contamination inside a UV hood. A unique box of sterile pipette tips was used for each sample. Between sample extractions, the area and pipettes were UV-radiated for 24 hours.

Gene regions were amplified via PCR with a 50 µl volume reaction using Qiagen Taq PCR Core Kit (Qiagen, Hilden, Germany) BRCA-1 and GHR forward and reverse primers^18,19^ at 0.5µM final concentration each, 200µM each dNTP, 1.25U Taq, 2.5 mM MgCl2, 3µl gDNA (100-1000 ng/µl). BRCA-1 PCR amplification conditions were 94°C 2 mins denaturation followed by 30 cycles: 94°C 30 s., 55.4°C 45 s., 72°C 45 s. and final at 72°C for 7 min. extension. GHR PCR amplification conditions were 94°C 2 mins denaturation followed by 30 cycles: 94°C 30 s., 54.8°C 45 s., 72°C 45 s. and final 72°C 7 min. final extension. The Cytb sequence was amplified using a 25 µl volume with Qiagen Taq PCR Core Kit (Qiagen: Valencia, CA) using PCR primers L14771 and H15149^25^ at 1.5µM each, 200µM each dNTP, 1.25U Taq, 1.5 mM MgCl2, and 4µl template DNA (100-1000 ng/µl.) PCR amplification conditions were 94°C 5 mins denaturation followed by 35 cycles: 94°C 1 min., 53.0°C 1 min., 72°C 1 min. and final at 72°C for 7 min. final extension.

We used microsatellite primers Trf254392, Trf276399, Trf5049, Trf52401, Trf58225, Trf661228, Trf677150, Trf71052, Mvis002, Mvis022, and Mvis072^21^; Mer049, and Mvis9700^22^; and Gg-14^23^. PCR protocols and multiplex reactions (MR) were established by^24^. For all single-plex reactions and MR 1, 4, and 5, PCR volumes consisted of 20-80 ng DNA, 1.6 pmol of each primer, and 5 uL of MeanGreen Master Mix (Empirical Bioscience, Grand Rapids, MI) in a 10 uL reaction volume. For reactions MR 2 and 3 PCR volumes consisted of 40-160 ng DNA, 1.6 pmol of each primer, and 10 uL of MeanGreen Master Mix in a 20 uL reaction volume. All polymerase chain reactions had an initial denaturation of 2 min at 95°C, followed by 39 cycles of 95°C for 10 s, primer-specific annealing temperature 53 - 56°C (Table 3) for 40 s, and a 30 s extension period at 72°C. A final extension time of 5 min at 70°C ended the reaction. The likelihood of contamination was minimized by running each individual four times at each locus and using a unique master mix for each locus for each sample. All PCR conditions were carried out under a UV hood, and similar to the extraction protocol the area and pipettes were UV radiated for 24 hours between PCR of the individuals, and a unique set of pipette tips were used for each individual. Sample Polymerase chain reaction products were analyzed on an ABI 3100 Genetic Analyzer using GeneScan Analysis 3.1.2, and genotypes were determined using Genotyper 4.0 software (all from Applied Biosystems, Foster City, California). Comparisons of ferret sample genotypes were performed using GenAlEx^25^. No discrepancies were found within individuals between PCR runs.

**Table 3:** Multiplex groupings and annealing temperatures of the 14 primers used for ferret clone verification.

| Multiplex Reaction (MR) | Primer | Annealing Temp (C ) |
| --- | --- | --- |
| 1 | Trf661228 | 54 |
|  | Mvis072 |  |
| 2 | Trf254392 | 54 |
|  | Trf71052 |  |
|  | Mvis9700 |  |
| 3 | Trf276399 | 54 |
|  | Mer049 |  |
|  | Mvis002 |  |
| 4 | Trf58225 | 56 |
|  | Trf5049 |  |
| 5 | Trf52401 | 56 |
|  | Gg-14 |  |
| Single-plex | Trf677150 | 54 |
| Single-plex | Mvis022 | 53 |

**Table 4.**
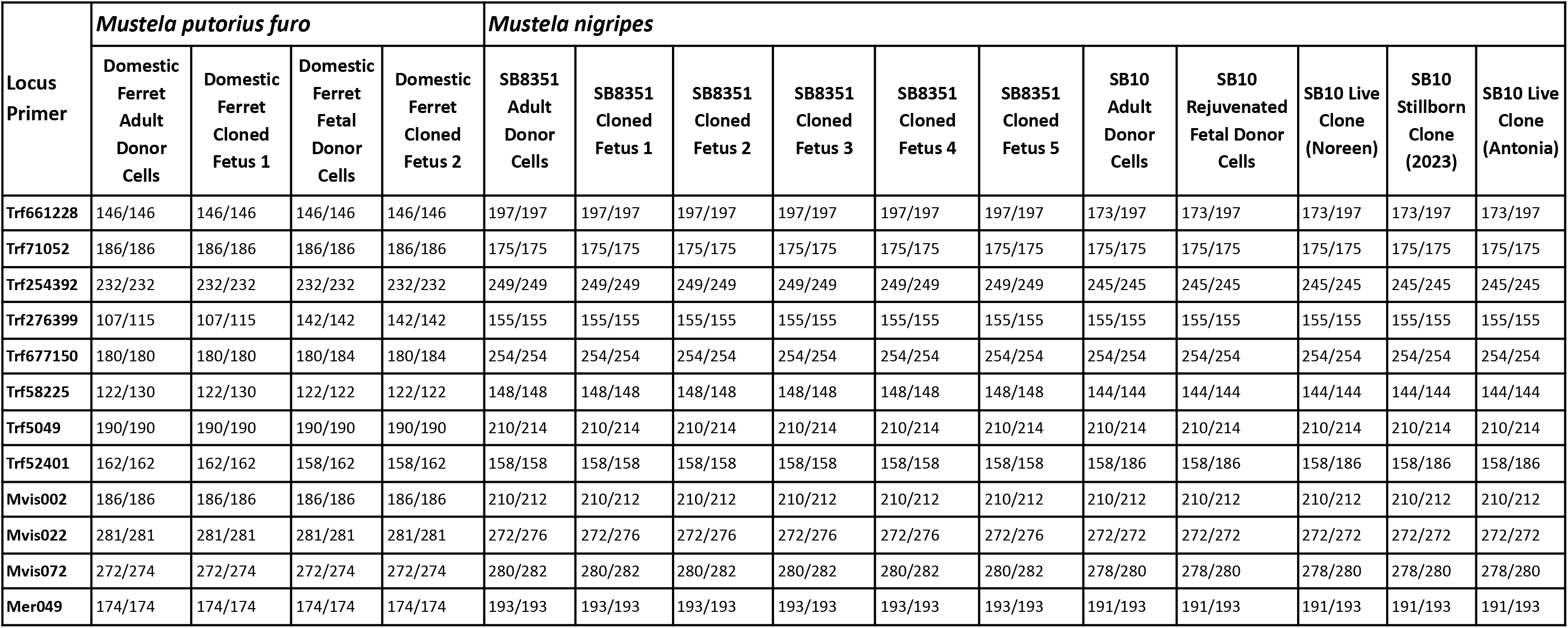
Microsatellite genotyping results for 12 successfully amplified loci. The Mvis9700 and Gg-14 loci failed to amplify for all samples and are excluded from the results.

**Table 5.** The percent of sequences of Cytb mtDNA that aligned to either domestic ferret or black-footed ferret. Numbers in parentheses are the number of TA TOPO clone sequences that aligned to a species divided by the total number sequenced for each sample.

| Sample | Nuclear Donor | Oocyte Donor / Maternal Genome | % *M. nigripes* mtDNA | % *M. p. furo* mtDNA | Nuclear DNA identity (BRCA-1) |
| --- | --- | --- | --- | --- | --- |
| Domestic Ferret Adult Donor Cells | *M. p. furo* | *M. p. furo* | 0 | 100% (10/10) | *M. p. furo* |
| Cloned Domestic Ferret Fetus 1 | *M. p. furo* | *M. p. furo* | 0 | 100% (10/10) | *M. p. furo* |
| Domestic Ferret Fetal Donor Cells | *M. p. furo* | *M. p. furo* | 0 | 100% (9/9) | *M. p. furo* |
| Cloned Domestic Ferret 2 | *M. p. furo* | *M. p. furo* | 0 | 100% (10/10) | *M. p. furo* |
| SB8351 Adult Donor Cells | *M. nigripes* | *M. nigripes* | 100% (8/8) | 0 | *M. nigripes* |
| SB8351 Cloned Fetus 1 | *M. nigripes* | *M. p. furo* | 0 | 100% (10/10) | *M. nigripes* |
| SB8351 Cloned Fetus 2 | *M. nigripes* | *M. p. furo* | 20% (2/10) | 80% (8/10) | *M. nigripes* |
| SB8351 Cloned Fetus 3 | *M. nigripes* | *M. p. furo* | 0 | 100% (10/10) | *M. nigripes* |
| SB8351 Cloned Fetus 4 | *M. nigripes* | *M. p. furo* | 0 | 100% (7/7) | *M. nigripes* |
| SB8351 Cloned Fetus 5 | *M. nigripes* | *M. p. furo* | 0 | 100% (9/9) | *M. nigripes* |
| SB10 Adult Donor Cells | *M. nigripes* | *M. nigripes* | 100% (6/6) | 0 | *M. nigripes* |
| SB10 Stillborn Clone (2020) | *M. nigripes* | *M. p. furo* | 11% (1/9) | 88% (8/9) | *M. nigripes* |
| SB10 Live Clone (Elizabeth Ann) | *M. nigripes* | *M. p. furo* | 20% (2/10) | 80% (8/10) | *M. nigripes* |
| SB10 Rejuvenated Fetal Donor Cells | *M. nigripes* | *M. p. furo* | 0 | 100% (11/11) | Not sequenced |
| SB10 Stillborn Clone (2023) | *M. nigripes* | *M. p. furo* | 0 | 100% (12/12) | Not sequenced |
| SB10 Live Clone (Noreen) | *M. nigripes* | *M. p. furo* | 0 | 100% (9/9) | Not sequenced |
| SB 10 Live Clone (Antonia) | *M. nigripes* | *M. p. furo* | 0 | 100% (12/12) | Not sequenced |

### Analysis of mitochondrial DNA of cloned black-footed ferrets by TA TOPO cloning and sequencing

Total DNA was extracted from cell lines derived from cloned fetuses of either black-footed ferret or domestic ferret. In addition, DNA was extracted from whole blood and buccal swabs collected from two clones born in 2020, the stillborn kit and Elizabeth Ann. DNA from all biosamples was extracted using the Gentra PureGene kit following sample-specific protocols (Qiagen Puregene Blood and Tissue Kit, Qiagen, Hilden, Germany). DNA extractions for three clones born in 2023, 1 stillborn, Noreen, and Antonia, were obtained from the microsatellite genotyping work.

Because the cloning process combines the cytoplasm of the nucleus donor and the oocyte donor, there is a chance that clones are heteroplasmic - containing a mix of both domestic ferret and black-footed ferret nuclear DNA. For some samples, the consensus sequences of the Cytb PCRs were a match for domestic ferrets, but the sequence chromatograms indicated low proportions of polymorphisms, potentially indicative of heteroplasmy. To investigate this we amplified the Cytb gene region again for use with a TOPO TA-Cloning kit (Invitrogen, Waltham, MA) to replicate and segregate different sequences originating from domestic ferret or black-footed ferret. Initial PCR products were inserted into the plasmid vector and the recombinant plasmids were transformed into E. coli. Up to 12 bacterial colonies that contained the plasmids were sequenced to determine SNP variants. Sanger sequencing performed at Eurofin Genomics (Eurofins Genomics, Louisville, KY)

We used the program Geneious Prime® 2023.2.1 (http://www.geneious.com) to trim, create consensus sequences from forward and reverse primers, and to align to known domestic and black-footed ferret sequences.

## Notes

### Competing Interest Statement

The authors have declared no competing interest.

